# Higher Temperatures Generically Favor Slower-Growing Bacterial Species in Multispecies Communities

**DOI:** 10.1101/689380

**Authors:** Simon Lax, Clare I. Abreu, Jeff Gore

## Abstract

Temperature is among the cardinal environmental variables which determine the composition and function of microbial communities. Many culture-independent studies have characterized communities that are affected by changing temperatures, either due to seasonal cycles^1-3^, long-term warming^4-6^, or latitudinal/elevational gradients^7-8^. However, a predictive understanding of how microbial communities respond to such changes in temperature is still lacking, partly because it is not obvious which aspects of microbial physiology determine whether a species should benefit from temperature alteration. Here, we incorporate how microbial growth rates change with temperature to a modified Lotka-Volterra competition model^9^, and predict that higher temperatures should generically favor slower-growing species in a bacterial community. We experimentally confirm this prediction in pairwise cocultures assembled from a diverse set of species, and we show that these changes to pairwise outcomes with temperature are also predictive of changing outcomes in three-species communities, suggesting our theory may propagate to more complex assemblages. Our results demonstrate that it is possible to predict how bacterial communities will shift with temperature knowing only the growth rates of the community members. These results provide a testable hypothesis for future studies of more complex, natural communities, and we hope that this work will help bridge the gap between ecological theory and the complex dynamics observed in metagenomic surveys.

---

Experimental microbial communities are normally incubated at a fixed temperature. We aimed to determine how changing this incubation temperature would affect the outcome of a microbial coculture in which the two species were known to stably coexist at our usual experimental temperature of 25°C. We focused on two naturally co-occurring species isolated from soil (Aci1 and Pan1), and followed a standard coculture methodology (see Methods) at three experimental temperatures: 16°C, 25°C, and 30°C. At each of these three temperatures, Aci1 is the faster growing species, and the difference in the growth rates of the two species increases alongside temperature (**Figure 1A**). Accordingly, we assumed that the slower-growing Pan1 would be favored by lowering the temperature and disfavored by raising the temperature, as its competitive ability would likely be hindered by a larger disparity in growth rate. Surprisingly, we observed the opposite, and found that Pan1 in fact becomes a *stronger* competitor at higher temperature, with the coculture outcome shifting from Aci1 dominance at 16°C (**Figure 1B**) to coexistence at 25°C (**Figure 1C**) and finally to Pan1 dominance at 30°C (**Figure 1D**).

**Figure 1:**
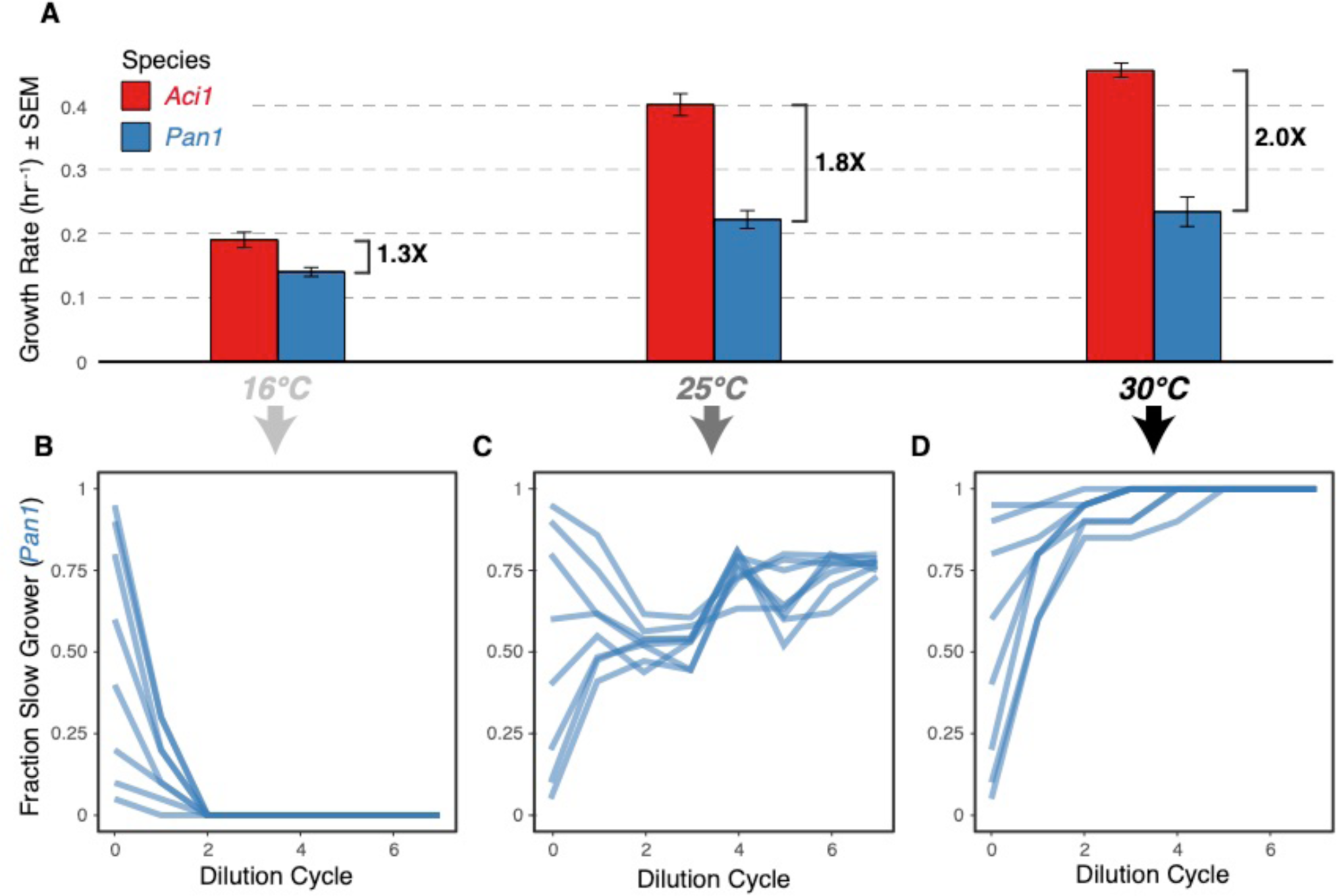
Increasing temperature favors the slower-growing bacterial species in a coculture, despite widening the difference in growth rates. This figure highlights the coculture outcomes of a faster-growing *Acinetobacter* species (Aci1) and a slower-growing *Pantoea* species (Pan1), both isolated from the same soil sample. (**A**) Aci1 is the faster grower regardless of temperature, and the difference in growth rates between the two species accelerates as temperature increases. Increasing temperature moves the equilibrium community state from competitive exclusion by Aci1 at 16°C (**D**) to coexistence at 25°C (**E**), and eventually to Pan1 dominance at 30°C (**F**), with the potentially counter-intuitive result that the slower growing species is favored by higher temperatures even when that change increases the gap in growth rates.

To explain this potentially counterintuitive result, we developed a model that expands on the work of Abreu *et al*.^**9**^, who used a modified version of the Lotka-Volterra competition model to explain how increasing mortality favors faster growing species. In addition to the growth rates, the Lotka-Volterra model requires knowledge of how the growth of the two species is inhibited by other cells of their own species as compared to the presence of cells of the competing species. This inhibition is traditionally captured by a parameter (α) that relates the strength of interspecific (between-species) competition to intraspecific (within-species) competition (**Figure 2A**). Competitive outcomes in the classic version of this model are determined entirely by these competition coefficients. However, many microbial communities experience mortality that is not driven by competition and which affects the entire community. Importantly, this is true of all laboratory cultures, where cells are removed from the community either continuously (as in a chemostat or turbidostat) or at discrete intervals (as in batch culture). It may also result from predation by bacterivores, or from physical removal, as in our gut microbiota. The formulation of the Lotka-Volterra model described above can therefore be made more realistic to microbial cocultures by the introduction of a community-wide mortality rate (*δ*). The introduction of this death rate to the model (**Figure 2B**) has an important effect: it makes the competitive outcome dependent on the growth rates as well as the competition coefficients, such that when the death rate is absorbed the α’s are reparametrized as

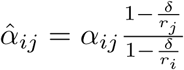

where *α*_*ij*_ is the inhibition of species *i* by species *j* without the death rate and 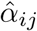 is the inhibition with the death rate. Adding mortality to the model favors the faster growing species^**9**^ by increasing *α*_*sf*_ and decreasing *α*_*fs*_. Visually, this change to the α’s of the two species can be represented as a 45-degree arrow through the phase space of the competitive outcomes (**Figure 2C**), pointing to the quadrant in which the faster grower wins. Mortality can reverse the competitive outcome if the slow grower would win without the mortality rate, passing first though a region of either coexistence or bistability. Importantly, the arrow is made longer by higher death rates and made shorter by higher growth rates. As bacterial growth rates are a function of temperature, this in turn introduces a temperature-dependence to the competition, and suggests that at any given death rate higher temperature should favor the slower growing species by lessening the favor conferred to the fast grower by the added mortality.

**Figure 2:**
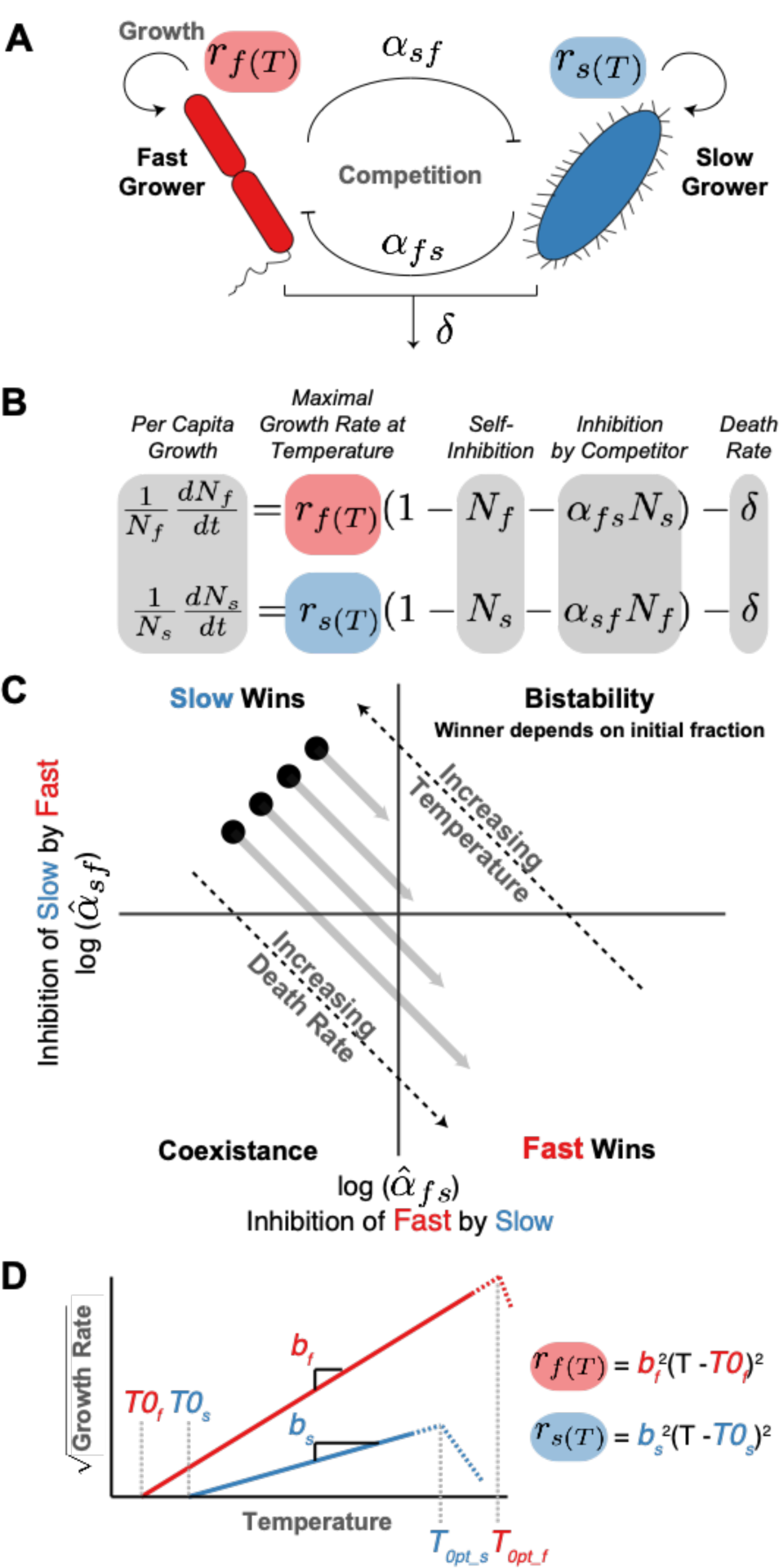
A simple model predicts that the slower-growing species in a coculture should generically be favored by increasing temperature. (**A**) The Lotka-Volterra competition models are parameterized by the growth rates of the two species and their competition coefficients (α), which relate between-species inhibition to within-species inhibition. A community-wide death rate (*δ*) can also be added to the model. (**B**) The full Lotka-Volterra competition equations, with added death rate. (**C**) With no death rate, competition outcomes are determined exclusively by the competition coefficients and do not depend on the growth rates (black points). The introduction of a death rate can alter the competitive outcome by effectively increasing the log(α) of the fast grower and decreasing the log(α) of the slow grower by the same amount, resulting in a 45-degree movement through phase space (light gray arrows). This arrow becomes longer as the death rate increases and shorter as temperature (and accordingly growth rates) increases, implying that for a given death rate, the slower-growing species should be favored by an increase in temperature. (**D**) The growth rates of microorganisms when sufficiently below T_Opt_ are a simple function of temperature that can be modeled with two parameters: the slope of the square-root of the growth rate against temperature (*b*) and the minimum growth temperature (*T0*).

To understand how temperature should influence the growth rate of bacterial species, we turned to the model of Ratkowsky *et al*.^**10**^. This phenomenological model predicts a linear relationship between temperature and the square root of a species’ growth rate, such that the growth rate of any bacterial species can be modeled (so long as it is sufficiently below the species’ optimum temperature, *T*_Opt_) as a function of two parameters: the slope of the presumed linear relationship (*b*), and the x-intercept of that line (*T0*) (**Figure 2D**). For any pair of species in which there is a consistent fast-grower (i.e. the faster grower has the lower *T0* and the higher *b*), plugging the Ratkowsky model into the competition model and taking the derivative of α_fs_ with respect to temperature reveals the surprising prediction that the slow grower is *always* favored by an increase in temperature. Remarkably, this is true even if the difference in growth rates between the two species increases with temperature. The prediction is largely generalizable to any temperature range in which there is a consistent faster-growing species and slower-growing species, even if their growth rate rankings flip far enough outside this temperature range (Supplementary Information). It is also generalizable to non-competitive interactions such as mutualism and parasitism (Supplementary Information). There are practical limits to this change to α: if the slow grower is already dominating at low temperatures then increasing temperature will not lead to a qualitative change in the outcome. Additionally, this model only holds when the temperature is below the species’ optimums, and growth rates increase alongside increases in temperature. Still, this theory suggests that it is possible to alter the competitive outcome by changing temperature, and to predict which species should benefit so long as the growth rates are known.

As a test of this theory, we chose a collection of 13 bacterial strains with variable growth rates (**Figure 3A**). This group comprised 6 strains from the ATCC culture collection and 7 naturally co-occurring strains isolated from soil. To fit the Ratkowsky model, we measured the growth rates of each strain at a minimum of four temperatures using a time to threshold approach (see Methods) (**Figure 3B, Supplementary Figure 1**). Both model parameters had a wide range, with *T0* ranging from −14°C to 4°C (mean = −3°C, SD = 5°C), and *b* ranging from 0.012 to 0.031 (mean = 0.024, SD = 0.005). Interestingly, these two values were highly correlated (ρ = 0.96, **Supplementary Figure 2**), suggesting that species which are capable of growing at lower temperatures (lower *T0*), are less able to increase their growth rates as temperature increases (lower *b*). This correlation has been previously reported in the literature^**11**^, although, to our knowledge, without any mechanistic explanation. It follows from the high correlation between *b* and *T0* that the curves representing the growth rate responses to temperature of different species (**Figure 3B**) are likely to intersect, such that which species we term the ‘fast grower’ and ‘slow grower’ may not be consistent across our range of temperatures. However, in 39 of the possible 78 pairs of species (50%) there was a consistent fast- and slow-grower across the range of experimental temperatures (16°C-30°C) (**Figure 3C**). We carried out 38 of these 39 pairwise cocultures, but did not coculture Pan1 and Pan2 because their colony morphologies are difficult to visually differentiate. For a subset of these cocultures, we varied the death rate as well as the temperature to explore how these two variables interact to shape competitive landscapes (Supplementary Information).

**Figure 3:**
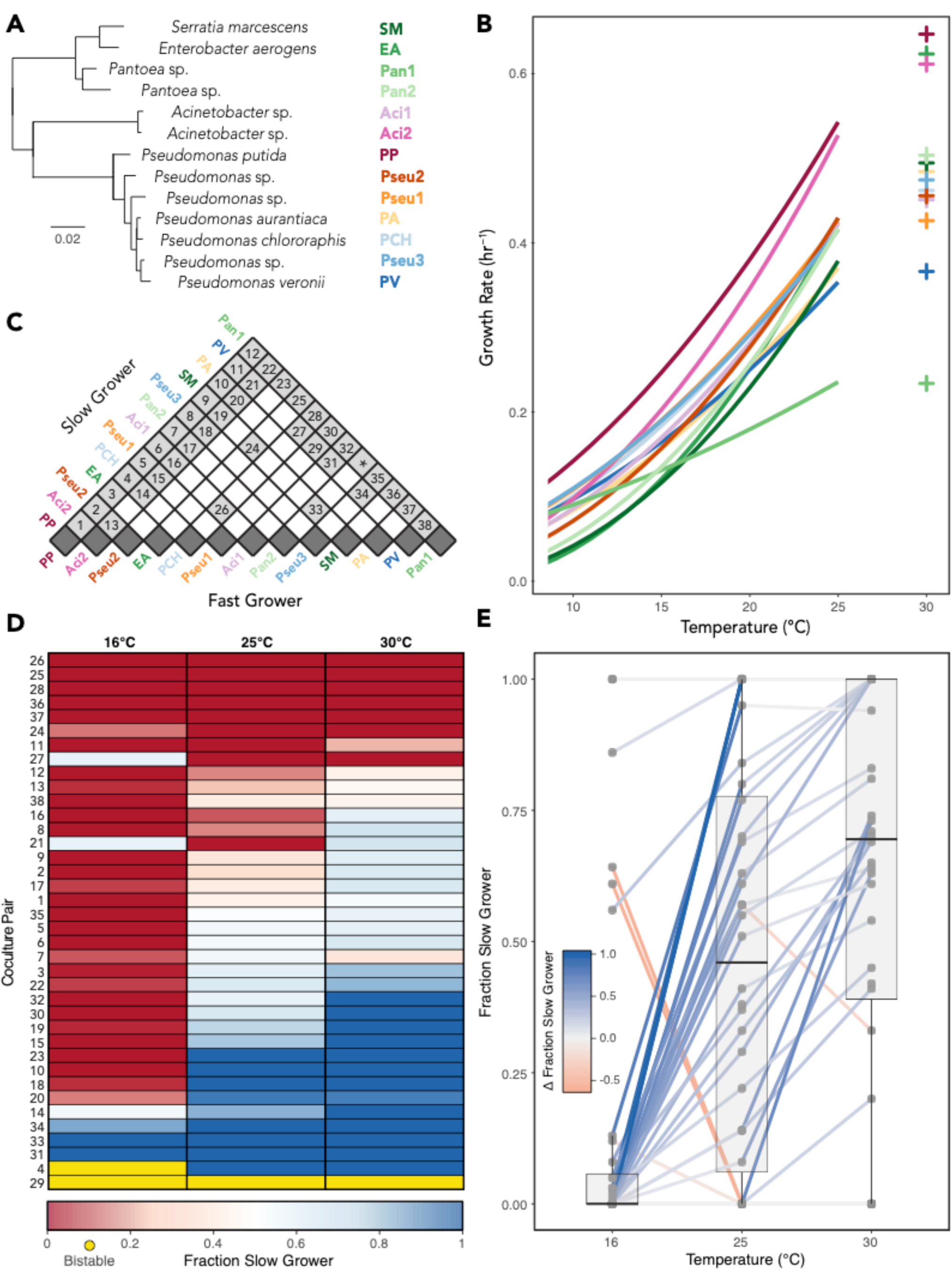
Theoretical predictions of which species should be favored by increasing temperature are validated in a wide array of experimental cocultures between a diverse set of species. (**A**) We tested our hypothesis that the slower-growing species should be favored by increasing temperature in model two-species communities drawn from 13 bacterial strains. Strains which are classified to the species level were obtained from the ATCC culture collection, and strains which are classified only to the genus level were isolated from a single soil sample and identified via sequencing of the 16S ribosomal subunit. Branch length of the phylogeny, based on full 16S sequences, corresponds to substitutions per base pair. (**B**) We measured the growth rates for each strain at a minimum of 4 temperatures in order to fit the Ratkowsky model for the range of 8°C to 25°C. 30°C growth rates, where the model may no longer hold, were measured directly. Line colors are matched to the label color for each strain in panel **A**. (**C**) Of the 78 possible species pairs, there were 39 pairs (50%) in which one strain was consistently the faster grower across the range of experimental temperatures (highlighted in gray). We cocultured 38 of these pairs at 16°C, 25°C, and 30°C following the experimental protocol of Figure 1. We did not coculture Pan1 and Pan2 because their colony morphologies are difficult to differentiate. (**D**) Heat map of coulure outcomes after a 7 day dilution cycle. Values are an average of at least 4 separate cocultures comprising 3 different initial strain ratios (90% fast grower, 50% fast grower, and 10% fast grower). Numbers designate species pairs as in **C**. (**E**) Boxplot of competitive outcomes at the three experimental temperatures. Black lines indicate the median, lower and upper box boundaries correspond to the first and third quartiles, and whiskers extend to the largest and smallest values within 1.5 times the inter-quartile range. Points indicate the outcomes of individual coculture pairs, and pairs are connected by lines, which are colored by the change in the mean equilibrium fast grower percentage. The two pairs in which we observe a bistable outcome are not included in the plot.

When these coculture outcomes are visualized as a heat map (**Figure 3D**), we observe a clear shift from fast-grower dominance at 16°C towards coexistence or slow-grower dominance in most species pairs. Plotting the changes in the slow-grower percentage for the two temperature shifts (**Figure 3E**) reveals that almost all transitions are in keeping with our theory. Of the 73 transitions that do not include a bistable outcome, 46 (63%) resulted in an increase in the slow-grower percentage in accordance with our theory, 23 (32%) led to no shift in the slow-grower percentage, and only 4 (5%) resulted in a decrease in the slow-grower percentage counter to our theory. In two of those four pairwise transitions not predicted by the model (EA/SM and Aci2/PV, both from 16°C to 25°C), the fast-grower dominated the community at 16°C when its initial fraction was 90% but coexistence was observed when its initial fraction was 50% or 10%, suggesting that the community may not have come to equilibrium within the 7 day experiment. In the third pair’s (PP/Pan2) transition from 25°C to 30°C, we always observed coexistence at both temperatures but with very high variance between replicates (0.1% - 0.8% slow grower at 25°C and 0% - 0.8% slow grower at 30°C), suggesting either experimental error or a high degree of stochasticity in this particular interaction. Finally, the fourth pair’s (PCH/PV) transition from 16°C to 25°C consistently showed a switch from coexistence to fast-grower dominance, suggesting some other temperature-dependent factor influenced the community in a direction counter to our theory.

Given the success of the model in predicting pairwise outcomes, we wanted to explore how temperature might impact more complex communities. Previous work in this group developed a simple predictive algorithm for inferring microbial community assembly from pairwise interactions^**12**^, which predicts that any species which is outcompeted in pairwise competition will not survive in any complex community that includes the other species in the pair. This implies that a change in temperature which shifts a pairwise interaction from competitive exclusion to coexistence could have broad implications for other species in the community, potentially resulting in cascading effects. The reverse is also possible: changes to temperature might shift a competitive outcome form coexistence to exclusion, decreasing the diversity of the community or allowing a species that was excluded by the newly outcompeted species to invade. We chose four trios of strains to test whether the changes we observed in the pairwise dynamics propagated to a three-species community. For each trio, we competed each pair of species in the trio from two initial species fractions and the full trio from four initial starting fractions (**Figure 4A**). We predicted that the community assembly rules should hold regardless of temperature and that increasing temperature should shift the equilibrium state away from the fastest grower and towards the slowest grower (**Figure 4A**).

**Figure 4:**
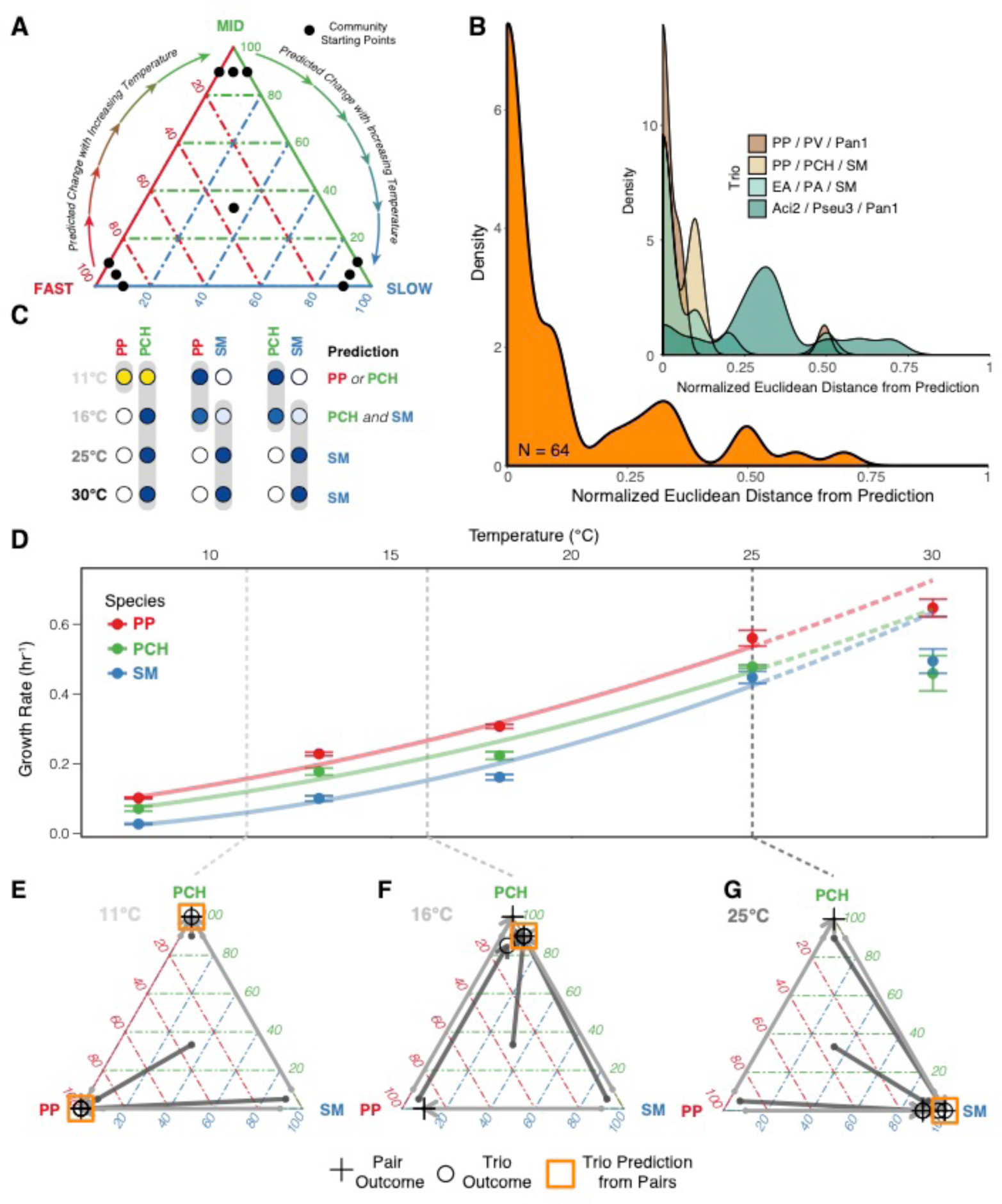
Shifts in pairwise competitive outcomes with temperature allow for prediction of the shifts observed in a three species community. (**A**) To test the predictive accuracy of pairwise dynamics for a three species community, we determined the equilibrium outcome of each pair in the trio from 2 initial starting points and the equilibrium outcome of the full trio from four starting points. When viewed as a ternary plot with the fastest species on the bottom-left and the slowest species on the bottom-right, we predict that increasing temperature should result in a clockwise movement of the equilibrium outcome. (**B**) We calculated the Euclidean distance (normalized to the maximum possible distance) between the predicted and observed equilibrium states. The outer density plot shows the distribution of distances across all four trios, while the inner plot splits the distribution by trio. (**C**) We used the assembly rules of Friedman**^12^** to estimate the equilibrium community state from the pairwise dynamics of each species in the trio. In this example, the pairwise dynamics predict a bistable outcome between PP and PCH at 11°C, coexistence between PCH and SM at 16°C, and dominance by SM at 25°C and 30°C. (**D**) This figure highlights the PP/PCH/SM trio, which has a consistent growth rate hierarchy regardless of temperature. (**E-G**) The experiment validates both the predicted clockwise movement about the ternary plot and the predictive accuracy of the assembly rules. Note that the 30°C outcome is not shown because it is identical to the 25°C treatment.

We found the community assembly rules were highly accurate: the standardized Euclidean distance from the prediction had a mean of 0.11, a SD of 0.17, and was 0 in more than half of the cases (32, 52%) (**Figure 4B**). Here, we focus on the PP/PCH/SM trio, in which the assembly rules predict a shift from bistable dynamics between PP and PCH at low temperature, to coexistence between PCH and SM at intermediate temperature, and ultimately to dominance by SM at high temperature (**Figure 4C**). In this trio, there is a consistent fast (PP) and slow (SM) grower across the full range of temperatures (**Figure 4D**), and the predictions from the pairwise dynamics are consistent with a movement in the equilibrium species fractions away from the fast grower and towards the slow grower as the temperature increases. This is in fact what we observed in our experiment: in almost all cases the equilibrium result was qualitatively the same as that predicted by the assembly rules. We also observed interesting dynamics in the PP/PV/Pan1 trio (**Supplementary Figure 3**). Here, PV was always excluded, but the species it was excluded by changed from PP to Pan1 as temperature increased. Based on the pairwise dynamics, there is possibly a temperature between 25°C and 30°C where PV should have been able to persist, highlighting how even very slight changes in temperature can alter the diversity of a microbial community.

All microbial communities are structured by interactions between their constituent species and between those species and the abiotic environment. As microbes compete for space and resources they have a number of tools at their disposal beyond the ability to grow faster^**13**^, including the production of secondary metabolites that are toxic to their competitors (antibiotics), contact-dependent inhibition, or antagonistic environmental alteration, for example through pH modification^**14**^. Temperature has the ability to influence each of these mechanisms, for example by varying the secondary metabolites produced by the community members^**15-16**^, manipulating the ability of community members to withstand the metabolites of other species^**17**^, or changing the pH range in which a species can grow^**18**^. Temperature may also play a role in determining the nutritional requirements of different species, potentially altering the nature of their ecological interactions and upsetting competitive hierarchies^**19,20**^. This complex set of interacting variables might suggest that predicting the effect of temperature on microbial competitive outcomes requires a potentially intractable knowledge of each species in the community and the interactions between them. However, we demonstrate here that we can obtain a great deal of predictive accuracy knowing nothing about the mechanisms which underpin those interactions and instead focusing exclusively on growth rates. This surprising simplicity may result from the density dependence of each of these competitive mechanisms: the greater the gap between a species’ growth rate and the death rate, the more the population of that species will be able to alter the environment in a manner favorable to itself. Even a very strong competitor will be rendered ineffectual if it is unable to reach a sufficient density. For example, a slower-growing strain which relies on antibiotic production as a competitive mechanism may not be able to produce a minimum inhibitory concentration if its growth rate is barely higher than the death rate.

Here, we demonstrate a potentially unifying predictive ability that only requires knowledge of a single variable: the maximal growth rate of each species. Although encouraging, these results are based on simple two- or three-species communities, drawn from a small species pool, in a tightly controlled lab environment. Still, this theory and preliminary experimental work provides a testable hypothesis for future studies of more complex, natural communities, and helps bridge the gap between ecological theory and the complex dynamics observed in metagenomic studies.

## MATERIALS & METHODS

### SPECIES & MEDIA

We used two sets of bacterial species in this study: seven naturally-co-occurring taxa isolated from soil and six strains ordered from the ATCC culture collection. The soil isolates were obtained by vortexing a small amount of soil taken from an urban park into PBS, followed by plating onto LB agar. Colonies were chosen based on the criteria that they were visually differentiable from all other strains in the study, and that they were capable of growing in the defined media described below. The taxonomic identity of the soil isolates was determined via sequencing of the V4-V5 16S hypervariable region, and the seven isolates were found to comprise representatives from three bacterial genera: *Acinetobacter* (Aci1 and Aci2), *Pantoea* (Pan1 and Pan2), and *Pseudomonas* (Pseu1, Pseu2, and Pseu3). The six species obtained from ATCC were *Enterobacter aerogenes* (EA, ATCC#13048), *Pseudomonas aurantiaca* (PA, ATCC#33663), *Pseudomonas chlororaphis* (PCH, ATCC#9446), *Pseudomonas putida* (PP, ATCC#12633), *Pseudomonas veronii* (PV, ATCC#700474) and *Serratia marcescens* (SM, ATCC#13880). All 13 strains are members of the bacterial class Gammaproteobacteria.

All coculture experiments in this study were carried out in S minimal medium supplemented with glucose (to a concentration of 0.2%) and ammonium chloride. The medium contained 100 mM sodium chloride, 5.7 mM dipotassium phosphate, 44.1 mM monopotassium phosphate, 5 mg/L cholesterol, 10 mM potassium citrate pH 6 (1 mM citric acid monohydrate, 10 mM tri-potassium citrate monohydrate), 3 mM calcium chloride, 3 mM magnesium sulfate, trace metals solution (0.05 mM disodium EDTA, 0.02 mM iron sulfate heptahydrate, 0.01 mM manganese chloride tetrahydrate, 0.01 mM zinc sulfate heptahydrate, 0.01 mM copper sulfate pentahydrate), 0.93 mM ammonium chloride, and 10 mM glucose.

### GROWTH RATE MODEL

To fit the Ratkowsky model for each strain, we calculated their growth rate at a minimum of four temperatures. We used a time to threshold approach to estimate growth rates, in which monocultures with known initial optical density (OD 600 nm) were spot checked every few hours. These growth rate experiments were carried out as follows: frozen stocks of the desired species were streaked out on a nutrient agar petri dish and, after incubation at room temperature for ∼48 hours, a single colony was picked into 5 mL of 1X LB broth and grown overnight. 35 uL of this LB culture was then inoculated into 5 mL of S medium and grown for ∼ 24 hours. The OD of the S medium culture was measured and the background OD (measured as the OD of the same volume of sterile S medium in the same type of 96 well plate) was subtracted in order to estimate population density. A log_10_ serial dilution of the monoculture was carried out on a 300 ul 96-well plate (Falcon) so that each strain was diluted to an OD of between 10^−1^ and 10^−6^ that of the overnight culture. The OD of each of these diluted cultures was checked periodically, the background OD was subtracted, and the growth rate was calculated as log(OD_*T*_/OD_*T*=0_)/*T* where OD_*T*=0_ is the initial OD of the diluted culture and OD_*T*_ is the OD at time *T* (measured as hours from initial time point). To make sure the cultures were still in their exponential phase of growth, growth rate was only calculated for measurements with OD_*T*_ < 0.15, and all growth rate estimates were based on a minimum of 5 measurements. This method of growth rate measurement implicitly incorporates lag time, as strains with a longer lag times will take longer to reach a given OD than another species with the same exponential growth rate but a shorter lag time.

### COCULTURE EXPERIMENTS

Frozen stocks of the competing species were streaked out on nutrient agar petri dishes and, after incubation at room temperature for ∼48 hours, a single colony of each species was picked into its own 50 mL Falcon tube containing 5 mL of 1X LB broth. Monocultures were grown overnight at room temperature, and 35 uL of this LB culture was then inoculated into 5 mL of S medium and grown for ∼ 24 hours at room temperature. The monocultures of each species were then OD-standardized, and the monocultures were mixed together with the desired proportions. In the two species experiments, the cocultures started from three initial community states: 90% fast grower / 10% slow grower, an equal split, and 10% fast grower / 90% slow grower. In the trio experiments, the competitions started from four initial community states: three 90%/5%/5% splits, each with a different species in the majority, and an even split of 33.3% of each species. All competition experiments were carried out in 300 uL 96 well plates (Falcon). The initial plate was made by adding 160 uL of S Media, 20 uL of 2% glucose, and 20 uL of a ^1^/_10_ dilution of the appropriate mixed cultures to each well, and was then incubated, wrapped in Parafilm and without shaking, for 24 hours at the desired temperature. Each day, for seven cycles, the previous day’s plate was serially diluted into new S Media so that each well held 180 uL of a ^1^/_100_ dilution of the mixed culture, and 20 uL of 2% glucose was added before incubation for another 24 hours. At the end of the competition cycle, the cultures were spot plated onto nutrient agar after dilution in phosphate-buffered saline, and colonies were counted by visual inspection to determine the equilibrium fraction of the species.

## CODE AND DATA AVAILABILITY

Access to the data is publicly available at https://doi.org/10.6084/m9.figshare.8285543.v1. All code for data analysis is available from the first author by request.

## ACKNOWLEDGEMENTS

We thank Anthony Ortiz for providing us with the bacterial soil isolates, and the members of the Gore lab for their suggestions and discussion.

## AUTHOR CONTRIBUTIONS

All authors designed the study. S.L. carried out the experiments with assistance from C.I.A. S.L. analyzed the data. C.I.A. analyzed the Lotka-Volterra and other models, and wrote Supplementary Notes VI – IX. S.L. wrote the manuscript and all authors edited and approved it.

## References

1. Gilbert, Jack A., et al. “Defining seasonal marine microbial community dynamics.” The ISME Journal 6.2 (2012): 298.

2. Fuhrman, Jed A., Jacob A. Cram, and David M. Needham. “Marine microbial community dynamics and their ecological interpretation.” Nature Reviews Microbiology 13.3 (2015): 133.

3. Ward, Christopher S., et al. “Annual community patterns are driven by seasonal switching between closely related marine bacteria.” The ISME journal 11.6 (2017): 1412

4. Barton, Andrew D., et al. “Anthropogenic climate change drives shift and shuffle in North Atlantic phytoplankton communities.” Proceedings of the National Academy of Sciences 113.11 (2016): 2964–2969.

5. Deslippe, Julie R., et al. “Long-term warming alters the composition of Arctic soil microbial communities.” FEMS microbiology ecology 82.2 (2012): 303–315.

6. Luo, Chengwei, et al. “Soil microbial community responses to a decade of warming as revealed by comparative metagenomics.” Appl. Environ. Microbiol. 80.5 (2014): 1777–1786.

7. Fuhrman, Jed A., et al. “A latitudinal diversity gradient in planktonic marine bacteria.” Proceedings of the National Academy of Sciences 105.22 (2008): 7774–7778.

8. Fierer, Noah, et al. “Microbes do not follow the elevational diversity patterns of plants and animals.” Ecology 92.4 (2011): 797–804.

9. Abreu, Clare I., et al. “Mortality causes universal changes in microbial community composition.” Nature communications10.1 (2019): 2120.

10. Ratkowsky, D. A., et al. “Relationship between temperature and growth rate of bacterial cultures.” Journal of bacteriology149.1 (1982): 1–5.

11. Rosso, L., J. R. Lobry, and J. P. Flandrois. “An unexpected correlation between cardinal temperatures of microbial growth highlighted by a new model.” Journal of Theoretical Biology162.4 (1993): 447–463.

12. Friedman, Jonathan, Logan M. Higgins, and Jeff Gore. “Community structure follows simple assembly rules in microbial microcosms.” Nature Ecology & Evolution 1.5 (2017): 0109.

13. Stubbendieck, Reed M., and Paul D. Straight. “Multifaceted interfaces of bacterial competition.” Journal of bacteriology198.16 (2016): 2145–2155.

14. Ratzke, Christoph, and Jeff Gore. “Modifying and reacting to the environmental pH can drive bacterial interactions.” PLoS biology 16.3 (2018): e2004248.

15. de Carvalho, Carla CCR, and Pedro Fernandes. “Production of metabolites as bacterial responses to the marine environment.” Marine Drugs 8.3 (2010): 705–727.

16. James, P. D. A., C. Edwards, and M. Dawson. “The effects of temperature, pH and growth rate on secondary metabolism in *Streptomyces thermoviolaceus* grown in a chemostat.” Microbiology 137.7 (1991): 1715–1720.

17. Sun, Wei, et al. “Mechanism and effect of temperature on variations in antibiotic resistance genes during anaerobic digestion of dairy manure.” Scientific reports 6 (2016): 30237.

18. Kim, C., and E. Ndegwa. “Influence of pH and temperature on growth characteristics of leading foodborne pathogens in a laboratory medium and select food beverages.” (2018).

19. Lewington-Pearce, Leah, et al. “Temperature-dependence of minimum resource requirements alters competitive hierarchies in phytoplankton.” Oikos (2019).

20. Hanke, Anna, et al. “Selective pressure of temperature on competition and cross-feeding within denitrifying and fermentative microbial communities.” Frontiers in Microbiology 6 (2016): 1461.

