## Supplementary Information for "Higher Temperatures Generically Favor Slower-Growing Bacterial Species in Multispecies Communities"

| | SECTION | *Page* |
|---|---|---|
| ***I*** | MEASUREMENT OF MAXIMAL GROWTH RATES | *2* |
| ***II*** | INTERACTION BETWEEN TEMPERATURE AND MORTALITY RATE | *3* |
| ***III*** | REMAINING OUTCOMES OF THREE-SPECIES MIXED CULTURES | *5* |
| ***IV*** | THE INFLUENCE OF TEMPERATURE ON NON-COMPETITIVE INTERACTIONS | *6* |
| ***V*** | CASES WHERE GROWTH RATE RANKS ARE NOT CONSISTENT ACROSS TEMPERATURES | *7* |
| ***VI*** | THE LOTKA-VOLTERRA PAIRWISE MODEL PREDICTS THAT INCREASING TEMPERATURE FAVORS THE SLOWER GROWER | *9* |
| ***VII*** | LINEAR RESOURCE CONCENTRATION MODEL WITH ONE RESOURCE PREDICTS THAT INCREASING TEMPERATURE FAVORS THE SLOWER GROWER | *11* |
| ***VIII*** | MONOD MODEL WITH ONE RESOURCE DOES NOT ALWAYS PREDICT THAT INCREASING TEMPERATURE FAVORS THE SLOWER GROWER | *13* |
| ***IX*** | MODELS WITH MORE THAN ONE RESOURCE DO NOT ALWAYS PREDICT THAT INCREASING TEMPERATURE FAVORS THE SLOWER GROWER | *15* |

**SUPPLEMENTARY NOTE I:** MEASUREMENT OF MAXIMAL GROWTH RATES

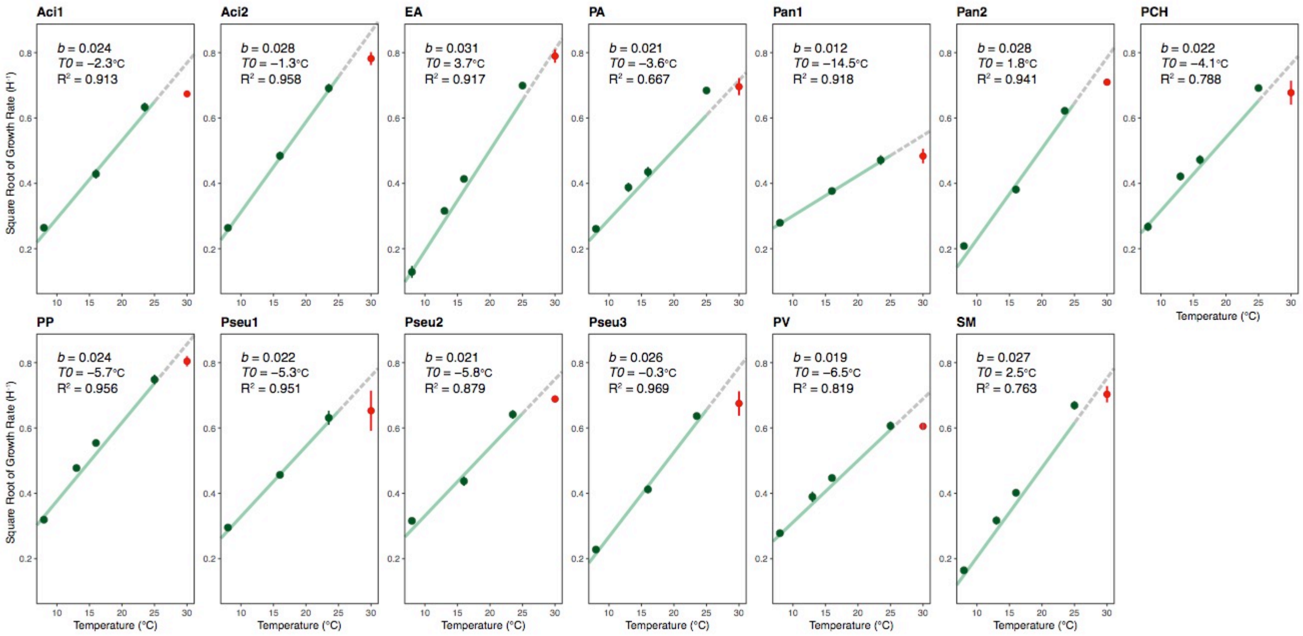


**Supplementary Figure 1: Fits of the Ratkowsky model to the 13 bacterial strains in this study.** The Ratkowsky model predicts a linear relationship between temperature and the square root of the growth rate. We used linear regression (light green line) on our growth rate estimates below 30°C (dark green circles, SEM indicated by bars) to estimate *b* (the slope of the regression) and *T0* (the x-intercept of the regression). The 30°C growth rates ± SEM are plotted in red. The $R^2$ of the regression is also provided.

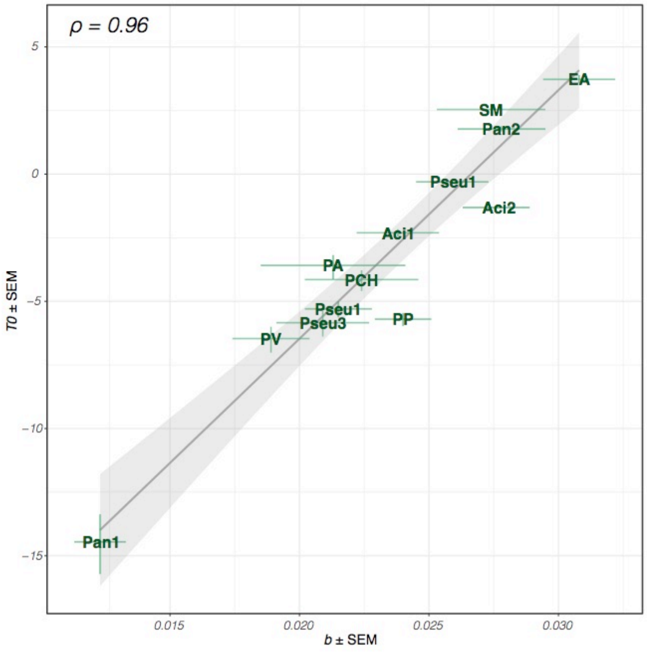


**Supplementary Figure 2: The two parameters of the Ratkowsky model are highly correlated.** Species are indicated by name and the gray line is a linear regression of the two parameters with the 95% confidence interval.

**SUPPLEMENTARY NOTE II:** INTERACTION BETWEEN TEMPERATURE AND MORTALITY RATE

To explore how the dilution factor and temperature interact to shape competitive outcomes, we used a factorial design to carry out competitions between two sets of species across four temperatures (11°C, 16°C, 25°C, and 30°C) and six dilution factors (a $\log_4$ serial dilution from $^1/_4$ to $^1/_{4096}$). Our theory predicts that increasing the dilution factor and increasing the temperature should move the competitive outcome in opposite directions along the 45 degree angle through phase space (**Figure 2D**), such that a given competition should never pass through both a bistable outcome and a coexistence outcome. Accordingly, we chose one species pair known to pass through a region of coexistence at our standard $^1/_{100}$ dilution factor (Aci1 & Pan1) and one pair known to pass through a region of bistability (PP & PCH).

In the Aci1/Pan1 competitions, the slower growing Pan1 always won at the lowest dilution rate ($^1/_4$) and at the highest temperature (30°C) (**Supplementary Figure 3A**). Increasing the dilution factor or decreasing the temperature always shifted the outcome to benefit the faster-growing Aci1, either to coexistence or competitive exclusion. This experimental outcome was qualitatively consistent with a model simulation of the competition, which used the measured *b* and *T0* values of the two strains and estimated α's to map the phase diagram of competitive outcomes (**Supplementary 3B**). In the PP/PCH competitions, we observed that the slower-growing PCH consistently won at the two highest temperatures (25°C and 30°C) and two lowest dilution factors, moving into a region of bistability when both temperature was low and the dilution factor was high (**Supplementary Figure 3C**). This outcome was also consistent with our model (**Supplementary Figure 3D**), although our theory predicts that regardless of the underlying α's there should always be a region right before collapse where the fast grower wins, first by competitive exclusion and then by default as the slow grower's growth rate drops below the death rate. This region of fast grower dominance may be quite small, and it is possible that the combination of temperature and dilution factor necessary to observe those dynamics was not included in the factorial design.

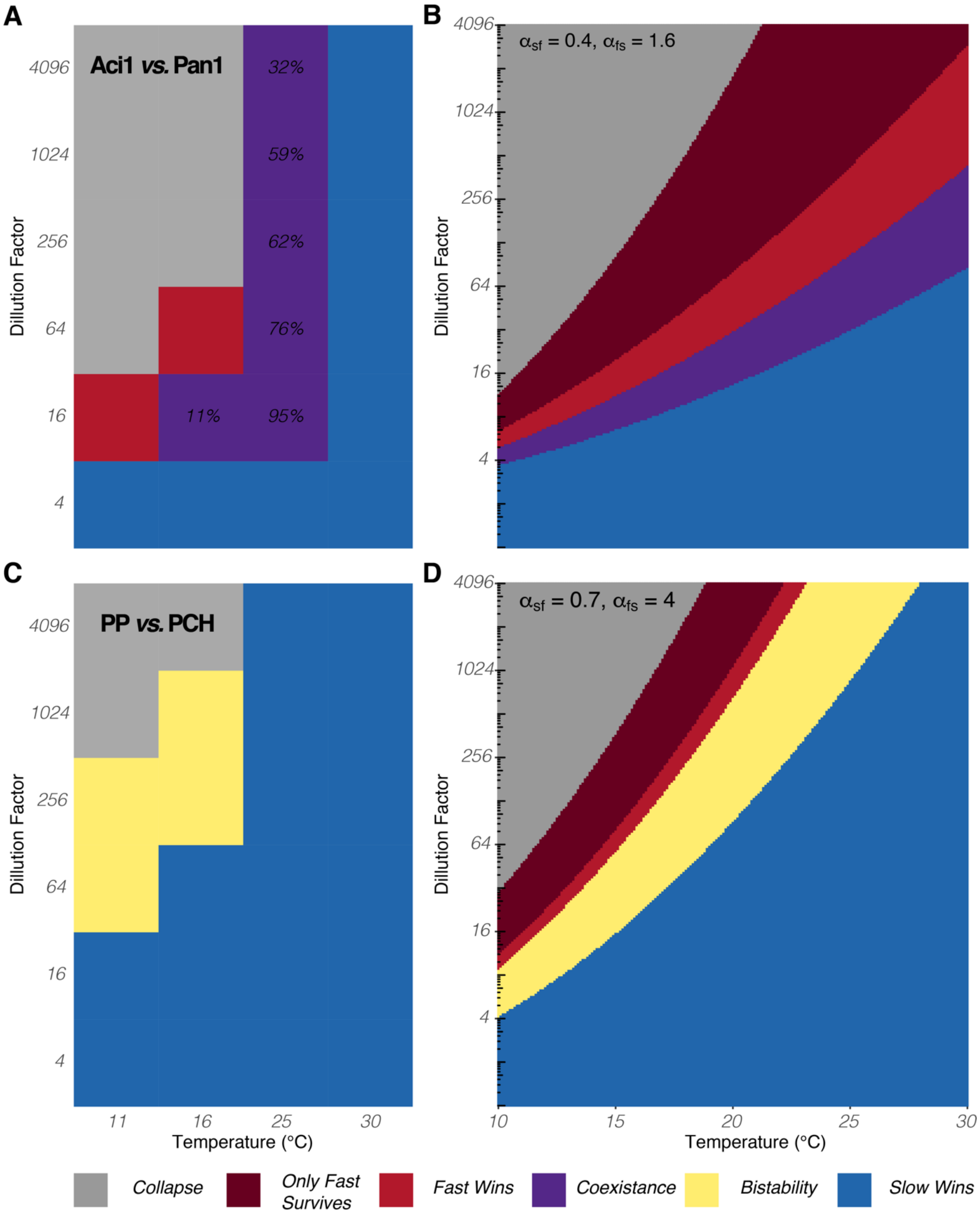


**Supplementary Figure 3: Phase diagrams of the interaction between temperature and dilution factor for two sets of coculture outcomes.** (**A**) We varied temperature (4 levels) and dilution factor (6 levels) in a factorial design to understand how these two variables interact to shape competitive landscapes. Here, we have the qualitative outcomes for the Aci1/Pan1 competition for each combination of variables. In cases of coexistence, the percentage of the final community comprised by the slow-grower is indicated. (**B**) Phase diagram of the model predictions for the Aci1/Pan1 competition, parameterized with the measured *b* and *T0* values, and with α's (indicated at top left) estimated to best match the experimental outcome. (**C**) & (**D**): As in **A** & **B**, but for the PP/PCH competition.

### **Supplementary Note III:** remaining outcomes of three-species mixed cultures

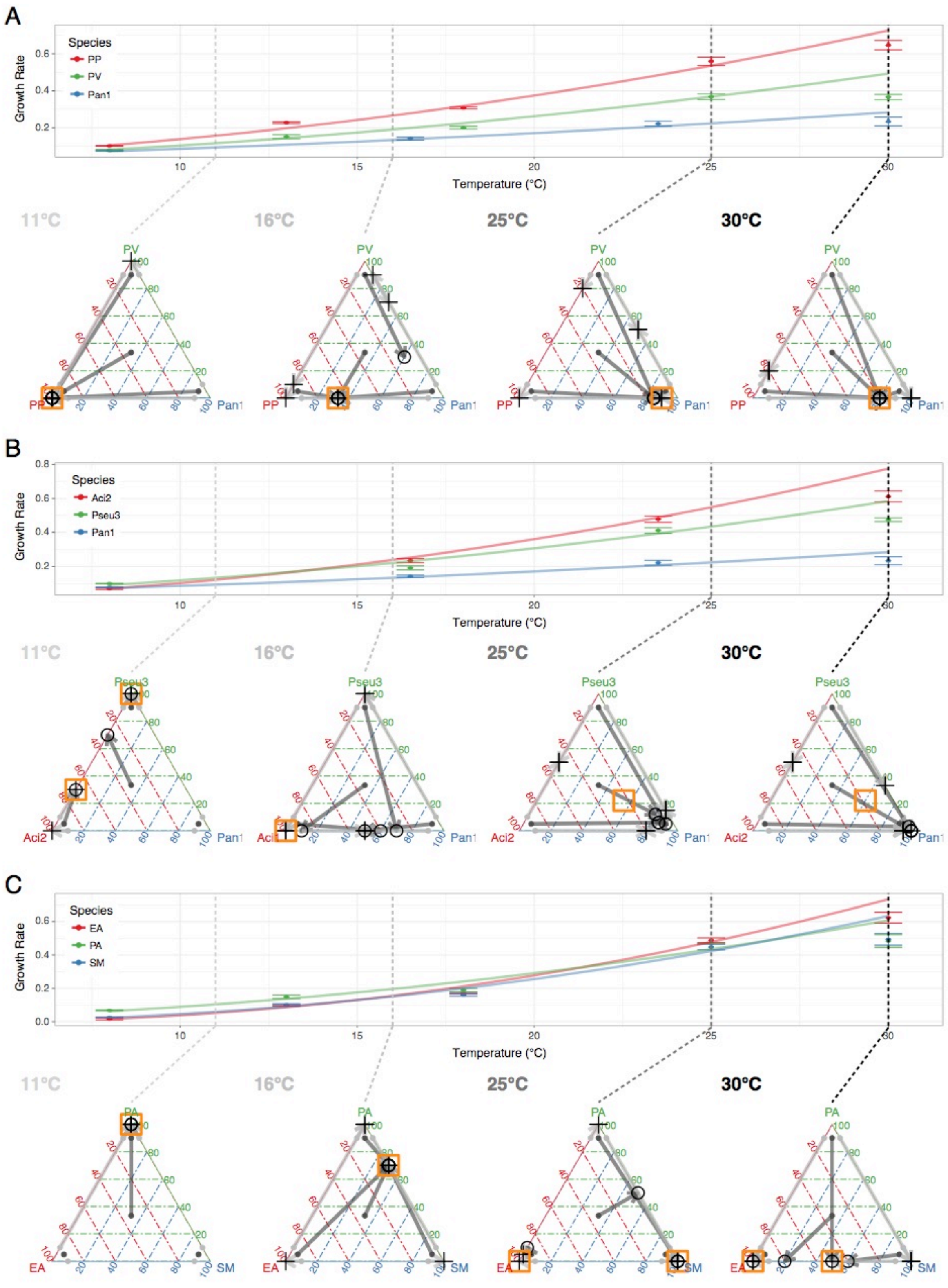


**Supplementary Figure 4: Coculture outcomes in the three three-species communities not plotted in Figure 4**. Growth rate plots are formatted as in **Figure 4D** and ternary plots are formatted as in **Figure 4E-G**.

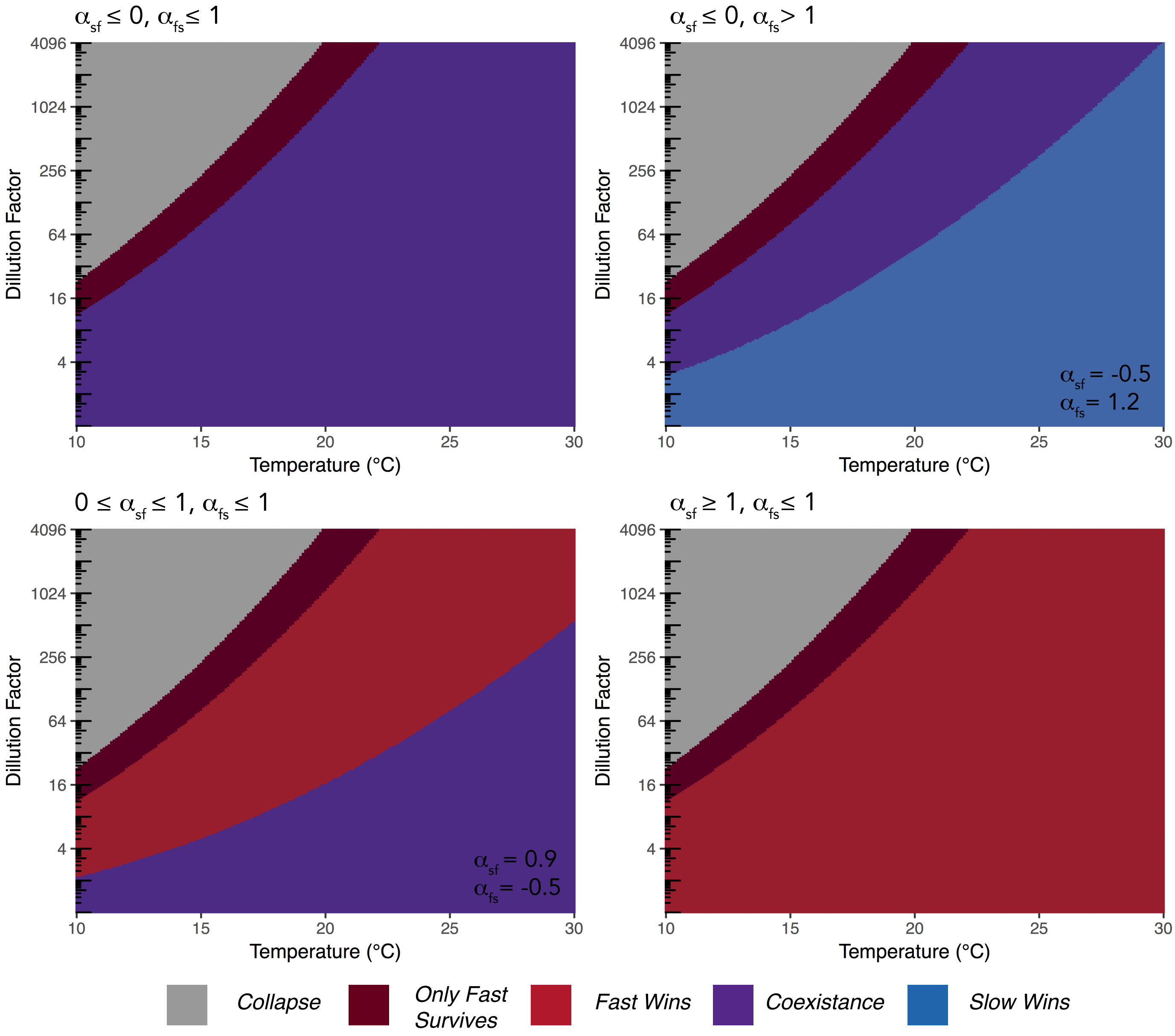


**Supplementary Figure 5: Predicted phase spaces of competitive outcomes with negative α's**. The model is based on a hypothetical faster-growing species with the parameters *b* = 0.0230 and *T0* = -5.7 and slower growing species with *b* = 0.0250 and *T0* = -4.1. In cases where there are multiple non-trivial qualitative outcomes, the specific α's are indicated at the bottom left of the plot. If no specific α's are indicated, the qualitative phase space is identical for all α's in the title range. In mutualist pairs, coexistence is always expected in any region of phase space where the slower-growing species is able to survive the imposed death rate, such that decreasing temperature can only move the outcome into a trivial fast-grower victory ($r_s < \delta < r_f$) or community collapse. Parasitic interactions, however, depend on the α's. If the fast grower assists the slow grower ($\alpha_{sf} < 0$) but the slow grower harms the fast grower ($\alpha_{fs} > 0$), we expect coexistence in all non-trivial regions of phase space ($r_s > \delta$) if $\alpha_{fs} < 1$ and a transition from coexistence to slow-grower dominance with increasing temperature if $\alpha_{fs} > 1$. If the slow grower assists the fast grower ($\alpha_{fs} < 0$) but is hindered by the fast grower ($\alpha_{sf} > 0$), we expect either consistent fast grower dominance if $\alpha_{sf} > 1$ or a transition from fast-grower dominance to coexistence with increasing temperature if $0 < \alpha_{sf} < 1$.

**SUPPLEMENTARY NOTE V:** CASES WHERE GROWTH RATE RANKS ARE NOT CONSISTENT ACROSS TEMPERATURES

Our theory always predicts a benefit to the slower-growing species with increasing temperature in cases where the growth rate rankings are consistent across all temperatures below $T_{Opt}$ (i.e. $T0_{Fast} < T0_{Slow}$ and $b_{Fast} > b_{Slow}$) (Supplementary Note VI). However, because of the high correlation between $T0$ and $b$, pairs of species that fit this criteria may be difficult to find, and there is no pair of species in this study that satisfies both those inequalities. Instead, we focus on pairs of species whose growth rate rankings do not cross within a defined temperature range, even if their growth rates cross at a lower or higher temperature. Although it is not generically true that the slower-growing species should be favored by increasing temperature in such cases, we can show that that prediction should hold true whenever the following inequality is met:

$$\frac{r_s'}{r_f'} > \frac{r_s(r_s-\delta)}{r_f(r_f-\delta)}$$

This inequality is always met in cases where the Ratkowsky growth curves do not cross, and is easily met generally so long as the growth rate of the slow grower is increasing with temperature at a rate that is not close to or equal to zero.

In cases where the growth rates intersect at a temperature below where both species are able to withstand the death rate, or in cases where the intersection occurs above the temperature range of interest, the prediction is straight forward: the slower-growing species is always favored by increasing temperature. In cases where the growth rates intersect at a temperature above which both species survive the death rate but below the temperature range of interest, the prediction in more complicated: there is an asymmetric effect where the slower-growing strain before the intercept benefits strongly from increasing temperature while the slower-growing strain after the intercept benefits comparably weakly (**Supplementary Figure 6**). This is because the benefits of an increasing growth rate are most profound near the temperature where a strain is first able to withstand the death rate, and relative yield is increasing the fastest.

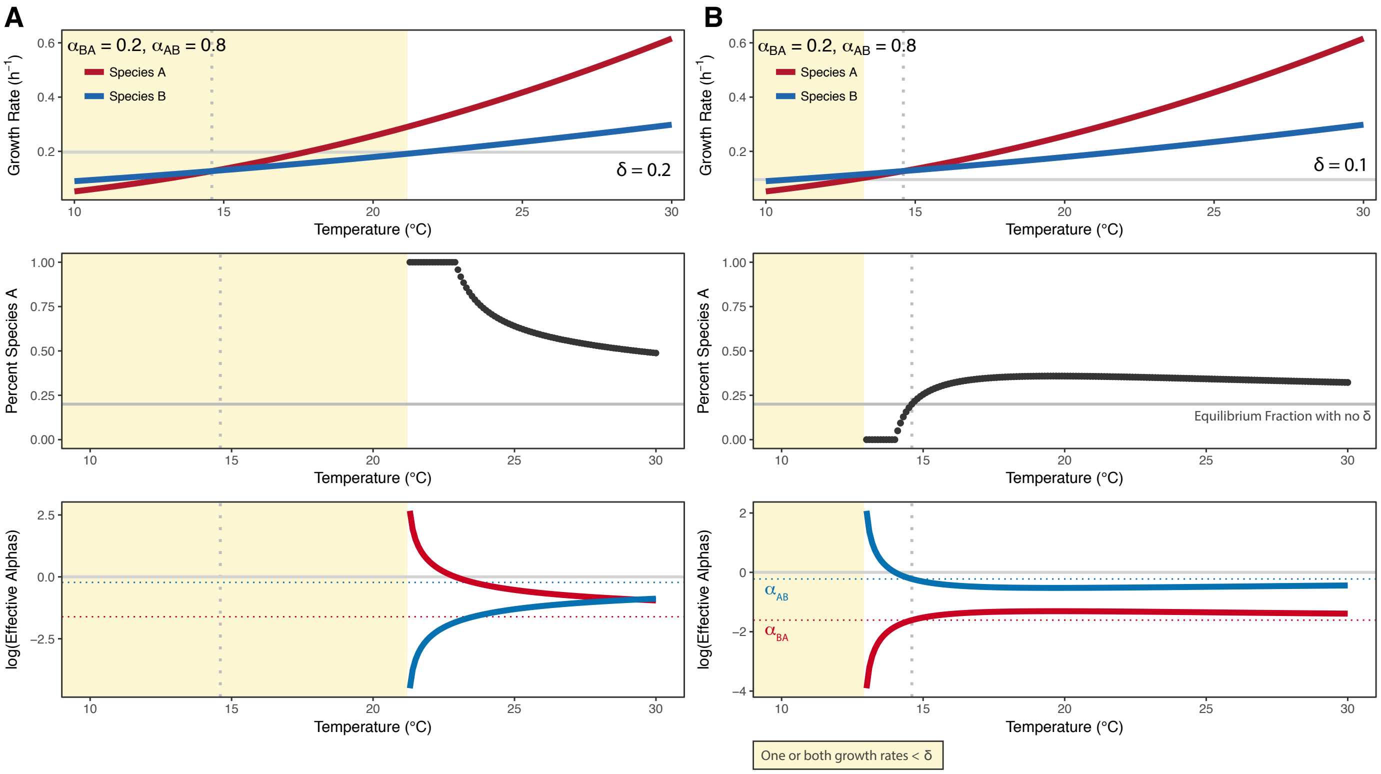


**Supplementary Figure 6: Coculture outcome predictions where growth rate responses to temperature cross.** This figure depicts the expected coculture outcomes for two hypothetical species with the following characteristics: $b_A$ = 0.028, $b_B$ = 0.012, $T0_A$ = 1.8°C, $T0_B$ = -14°C, $\alpha_{BA}$ = 0.2, and $\alpha_{AB}$ = 0.8. At a death rate of 0.2 per hour (**A**), the growth rates cross at a temperature below where both species are able to withstand the death rate (top), and species B, which is always the slower grower in this non-trivial temperature range, consistently increases its equilibrium fraction with increasing temperature (middle), as its α against A increases (bottom). At a lower death rate of 0.1 per hour (**B**), where the growth rates cross at a non-trivial temperature, species A increases its equilibrium fraction with temperature in the region where it is the slower grower, but continues to do so for a short range of temperatures after it becomes the faster grower before the direction of change flips to favor the slower grower again. Note that in this case the derivative of the change in species fraction with respect to temperature is much higher for the region before the growth rates cross, where the effect of the death rate is more pronounced, than it is after the line cross, where the death rate becomes increasingly dwarfed by the growth rates.

**SUPPLEMENTARY NOTE VI:** LOTKA-VOLTERRA PAIRWISE MODEL PREDICTS THAT INCREASING TEMPERATURE FAVORS THE SLOWER GROWER.

In this paper, we have found that increasing temperature generally favors slower-growing species in coculture experiments. To explain this result mathematically, we have relied on the predictions of the two-species Lotka-Volterra (LV) interspecific competition model:

$$\frac{\dot{N}_i}{N_i} = r_i\left(1 - N_i - \alpha_{ij}N_j\right) \tag{1}$$

where $N_i$ represents the abundance of species $i$ normalized by its carrying capacity, $r_i$ its maximal growth rate, and $\alpha_{ij}$ is a dimensionless competition coefficient quantifying the inhibition of species $i$ by species $j$. The outcomes of the model are completely determined by the competition coefficients: when both $\alpha_{ij} < 1, \alpha_{ji} < 1$, both species coexist; when $\alpha_{ij} < 1$ but $\alpha_{ji} > 1$, species $i$ drives species $j$ to extinction (and vice versa); when both $\alpha_{ij} > 1, \alpha_{ji} > 1$, the result is bistability, in which both species can drive each other extinct, and the winner depends on the initial relative fraction. Thus, the model does not depend on the growth rates—a strongly competing slow grower can drive a fast grower extinct.

Many microbial communities experience mortality that is not driven by competition and which affects the entire community. Importantly, this is true of all laboratory cultures, where cells are removed from the community either continuously (as in a chemostat or turbidostat) or at discrete intervals (as in batch culture). It may also result from predation by bacterivores, or, in the case of our gut microbiota, from waste passing through. Equation (1) can therefore be made more realistic by the introduction of a community-wide mortality rate ($\delta$):

$$\frac{\dot{N}_i}{N_i} = r_i\left(1 - N_i - \alpha_{ij}N_j\right) - \delta \tag{2}$$

Equation (2) can then be re-parameterized back into is original form, such that the outcome is determined completely by the re-parameterized competition coefficients $\hat{\alpha}_{ij}$:

$$\frac{\dot{\hat{N}}_i}{\hat{N}_i} = \hat{r}_i\left(1 - \hat{N}_i - \hat{\alpha}_{ij}\hat{N}_j\right) \tag{3}$$

$$\hat{\alpha}_{ij} = \alpha_{ij}\frac{(1 - \frac{\delta}{r_j})}{(1 - \frac{\delta}{r_i})} \tag{4}$$

Since $\hat{\alpha}_{ij}$ is a function of growth and death rates, the outcome will shift along with these rates. For example, the faster grower is favored by an increasing death rate. An increasing temperature, on the other hand, will affect growth rather than death. If we assume that maximal growth rates $r_i(T)$ are the only parameters affected by changes in temperature, we can take the derivative of $\hat{\alpha}_{ij}$ with respect to temperature to see which species will benefit from an increase in temperature:

$$\frac{\partial}{\partial T}\hat{\alpha}_{fs} = \alpha_{fs}\frac{\partial}{\partial T}\left(\frac{1 - \frac{\delta}{r_s(T)}}{1 - \frac{\delta}{r_f(T)}}\right) \tag{5}$$

Here was have used indices $f$ and $s$ to denote fast and slow grower, respectively. We must choose a model for $r(T)$ in order to continue. As discussed in the main text, the Ratkowsky model is consistently the best fit for the data:

$$r(T) = b^2(T - T_o)^2 \tag{6}$$

where $b$ and $T_o$ must be determined by fitting data for a particular species. Plugging this form into Equation 5, we find:

$$\frac{\partial}{\partial T}\hat{\alpha}_{fs} = \alpha_{fs}\frac{2\delta}{b_s^2(T - T_{os})^3(1 - \frac{\delta}{b_f^2\left(T - T_{of}\right)^2})} - \frac{2\delta(1 - \frac{\delta}{b_s^2(T - T_{os})^2})}{b_f^2\left(T - T_{of}\right)^3\left(1 - \frac{\delta}{b_f^2\left(T - T_{of}\right)^2}\right)^2} \tag{7}$$

For the slow-grower to be favored, the above term should be greater than zero, because this would mean that inhibition of the fast grower by the slow grower increases with temperature (and inhibition of the slow grower by the fast grower decreases):

$$\frac{2\delta}{b_s^2(T - T_{os})^3(1 - \frac{\delta}{b_f^2\left(T - T_{of}\right)^2})} > \frac{2\delta(1 - \frac{\delta}{b_s^2(T - T_{os})^2})}{b_f^2\left(T - T_{of}\right)^3\left(1 - \frac{\delta}{b_f^2\left(T - T_{of}\right)^2}\right)^2} \tag{8}$$

We can simplify this expression by plugging in $r_f$ and $r_s$ where they appear:

$$\frac{1}{r_s(T - T_{os})(1 - \frac{\delta}{r_f})} > \frac{1 - \frac{\delta}{r_s}}{r_f\left(T - T_{of}\right)\left(1 - \frac{\delta}{r_f}\right)^2} \tag{9}$$

Ultimately, the expression simplifies to a simple relation shows that the slow grower is always favored as temperature increases when the growth curves of the two species do not cross:

$$\frac{\left(T - T_{of}\right)}{(T - T_{os})} > \frac{r_s - \delta}{r_f - \delta} \tag{10}$$

When the growth curves do not cross, we can assume that $T_{os} > T_{of}$ and $r_f > r_s$. These assumptions make the left side of the inequality greater than one, while the right side is less than one. Thus, the inequality is always true for a competition between a consistent slow grower and a consistent fast grower, and the LV model predicts that an increasing temperature will always favor a slower grower, provided that the slower grower does not become relatively faster at high temperature.

The assumption of a community-wide death rate might be unrealistic—for example, a bacteriovore might preferentially prey on one species over the other. Accordingly, we can generalize our prediction to a scenario in which the death rate experienced by the slow grower ($\delta_s$) is not equal to that experienced by the fast grower ($\delta_f$):

$$\frac{\dot{N}_f}{N_f} = r_f\left(1 - N_f - \alpha_{fs}N_s\right) - \delta_f \tag{11}$$

$$\frac{\dot{N}_s}{N_s} = r_s\left(1 - N_s - \alpha_{sf}N_f\right) - \delta_s \tag{12}$$

In this case, Equation 4 becomes slightly modified:

$$\hat{\alpha}_{fs} = \alpha_{fs}\frac{(1 - \frac{r_s}{\delta_s})}{(1 - \frac{r_f}{\delta_f})} \tag{13}$$

Taking the derivative with respect to temperature and setting the result to be greater than zero yields an expression similar to Equation 10:

$$\frac{T - T_{of}}{T - T_{os}} > \left(\frac{r_s - \delta_s}{r_f - \delta_f}\right)\left(\frac{\delta_f}{\delta_s}\right) \tag{14}$$

The inequality is still always true if $\delta_s \geq \delta_f$. Thus, a slower grower will always be favored by an increase in temperature, assuming that the growth curves do not cross, and that it experiences a death rate equal to or higher than that experienced by the fast grower.

### **SUPPLEMENTARY NOTE VII:** LINEAR RESOURCE CONCENTRATION (LRC) MODEL WITH ONE RESOURCE PREDICTS THAT INCREASING TEMPERATURE FAVORS THE SLOWER GROWER.

The simplicity of the Lotka-Volterra model makes it a useful null model for experimental ecologists. Some may characterize it as too simplistic, however, because it quantifies interactions between species without specifying their mechanism. In resource-explicit models, species' interactions are mediated by resource consumption. The simplest form of such a model involves two species $N_1$ and $N_2$ competing for one resource $c$:

$$\frac{\dot{N}_1}{N_1} = r_1 c - \delta \tag{15}$$

$$\frac{\dot{N}_2}{N_2} = r_2 c - \delta \tag{16}$$

$$\dot{c} = \delta(c_o - c) - r_1 c N_1 - r_2 c N_2 \tag{17}$$

As in the LV model, $r$ represents growth rate. $\delta$ represents both the community-wide death rate as well as the influx rate of fresh nutrients, at concentration $c_o$, as would happen in a chemostat where nutrients flow in and cells are pumped out at the same rate. The species that can consume the resources at the highest per-capita rate will drive the other species extinct by decreasing the resource concentration to a lower equilibrium level $c^*$, which can be found by setting Equations 12 and 13 to zero:

$$c_1^* = \frac{\delta}{r_1} \tag{18}$$

$$c_2^* = \frac{\delta}{r_2} \tag{19}$$

A lower value of $c^*$ corresponds to a higher maximal per-capita growth rate $r$; species 1 excludes species 2 if it has a higher maximal growth rate and is therefore the faster grower:

$$c_1^* < c_2^* \tag{20}$$

$$\frac{\delta}{r_1} < \frac{\delta}{r_2} \tag{21}$$

$$r_1 > r_2 \tag{22}$$

Now we can ask whether an increase in temperature will favor the slower grower $N_2$. In order for this to happen, the difference between $c_1^*$ and $c_2^*$ must decrease such that $c_1^* - c_2^*$ becomes less negative:

$$\frac{\partial}{\partial T}(c_1^* - c_2^*) > 0 \tag{23}$$

Again we will assume that growth rate depends on temperature according to the Ratkowsky model:

$$\frac{\partial}{\partial T}\left(\frac{\delta}{b_s^2(T - T_{os})^2} - \frac{\delta}{b_f^2\left(T - T_{of}\right)^2}\right) > 0 \tag{24}$$

We have changed indices ($1 \rightarrow f, 2 \rightarrow s$) to denote that $N_1$ is the fast grower and $N_2$ is the slow grower. Taking the derivative yields nearly the same expression as in the analysis of the Lotka-Volterra model (Equation 10):

$$\frac{T - T_{of}}{T - T_{os}} > \frac{r_s}{r_f} \tag{25}$$

The prediction that an increase in temperature favors the slower grower is therefore always true in the LRC model with one resource. Again, we assume that the growth curves of the two species do no cross, which would cause the slower grower to become relatively faster at a higher temperature.

Again, we can generalize this prediction to the case of species-specific death rates. Equation 25, the condition for the slow grower to be favored by increasing temperature, is modified slightly:

$$\frac{T - T_{of}}{T - T_{os}} > \left(\frac{r_s}{r_f}\right)\left(\frac{\delta_f}{\delta_s}\right) \tag{25}$$

As in the analysis of the LV model (Equation 14) the inequality is still always true if $\delta_s \geq \delta_f$. In both the LV model and the LRC model with one resource, a slower grower will always be favored by an increase in temperature, assuming that the growth curves do not cross, and that it experiences a death rate equal to or higher than that experienced by the fast grower.

**SUPPLEMENTARY NOTE VIII:** MONOD MODEL WITH ONE RESOURCE DOES NOT ALWAYS PREDICT THAT INCREASING TEMPERATURE FAVORS THE SLOWER GROWER.

In the LRC model, per-capita growth rate increases linearly with resource concentration. This assumption is ideally suited to low resource concentrations, but becomes unrealistic at high concentrations. For this reason, the Monod model might better describe dynamics at high resource concentration, because the per-capita growth saturates to $r$ as $c$ grows large:

$$\frac{\dot{N}_1}{N_1} = \frac{r_1 c}{K_1 + c} - \delta \tag{26}$$

$$\frac{\dot{N}_2}{N_2} = \frac{r_2 c}{K_2 + c} - \delta \tag{27}$$

$$\dot{c} = \delta(c_o - c) - \frac{r_1 c N_1}{K_1 + c} - \frac{r_2 c N_2}{K_2 + c} \tag{28}$$

Here, $K$ is the half-maximal constant, or the resource concentration at which a species grows at half its maximum rate. The dominant species can again be found by setting the first two equations to zero and determining which species can drive the resource to a lower equilibrium concentration $c^*$:

$$c_1^* = \frac{\delta K_1}{r_1 - \delta} \tag{29}$$

$$c_2^* = \frac{\delta K_2}{r_2 - \delta} \tag{30}$$

In contrast to the LRC model, if species $1$ has a faster growth rate, it will not necessarily drive species $2$ extinct. We can see this by setting $c_f^*$ to be less than $c_s^*$:

$$\frac{\delta K_f}{r_f - \delta} < \frac{\delta K_s}{r_s - \delta} \tag{31}$$

$$\frac{r_f - \delta}{r_s - \delta} < \frac{K_f}{K_s} \tag{32}$$

While quantity on the left side of the inequality is always greater than one, the quantity on the right can span a wide range, and possibly be greater than the quantity on the left. A slow grower with a small half-maximal constant might outcompete a fast grower with a large half-maximal constant, because the slow grower might be relatively faster at low resource concentration. This tradeoff between $K$ and $r$ can be visualized by plotting per-capita growth vs. resource concentration for both species (**Supplementary Figure 7**).

To find the condition in which an increasing temperature favors the slower grower, we again set the derivative of the difference of equilibrium concentrations to be positive:

$$\frac{\partial}{\partial T}\left(c_f^* - c_s^*\right) > 0 \tag{33}$$

$$\frac{\partial}{\partial T}\left(\frac{\delta K_f}{b_f^2\left(T - T_{of}\right)^2 - \delta} - \frac{\delta K_s}{b_s^2(T - T_{os})^2 - \delta}\right) > 0 \tag{34}$$

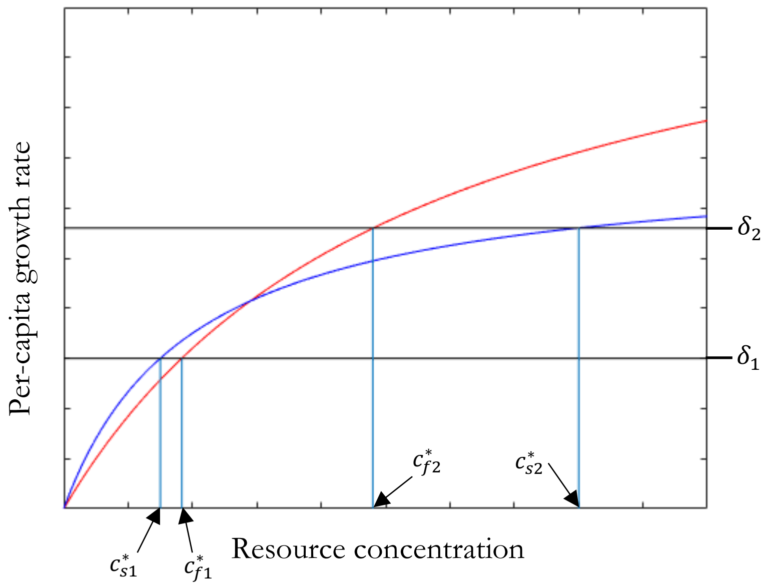


**Supplementary Figure 7**: A tradeoff between $K$ and $r$ can be seen as lines that cross on a plot of per-capita growth vs. resource concentration. At low death rate $\delta_1$, the blue species has a lower $c^*$ and thus dominates, despite having a lower maximal rate $r$. This is due to the fact that is also has a lower half maximal constant $K$. A higher death rate $\delta_2$ favors the faster grower. An increasing temperature is not guaranteed to favor the slower grower at either of these death rates, however. Such a prediction is guaranteed only when there is no tradeoff between $K$ and $r$ (see discussion below).

$$\frac{K_s b_s^2 (T - T_{os})}{(b_s^2 (T - T_{os})^2 - \delta)^2} > \frac{K_f b_f^2 (T - T_{of})}{(b_f^2 (T - T_{of})^2 - \delta)^2} \tag{35}$$

$$\left(\frac{r_f - \delta}{r_s - \delta}\right)^2 > \left(\frac{T - T_{os}}{T - T_{of}}\right)\left(\frac{K_f}{K_s}\right)\left(\frac{r_f}{r_s}\right) \tag{36}$$

The left side of this inequality is greater than one. The first term on the right side is less than one, and the second term is less than the square root of left side if there is no tradeoff between $K$ and $r$ (Equation 32, **Supplementary Figure 7**). Additionally, the third term on the right is also less than the square root of the left side. Hence, this inequality is always true if there is no tradeoff between $K$ and $r$, and in such a case, the slow grower will always be favored by an increase in temperature. The Monod model therefore makes a subtler prediction than the LRC and LV models that increasing temperature favors the slower grower—this is true only if the slow grower does not become relatively faster at either high temperature or low resource concentration.

**SUPPLEMENTARY NOTE IX:** MODELS WITH MORE THAN ONE RESOURCE DO NOT ALWAYS PREDICT THAT INCREASING TEMPERATURE FAVORS THE SLOWER GROWER.

The LRC and Monod models can be extended to include any number of species and resources. We chose to focus on two species for the purpose of comparing a slow grower and a fast grower. Whether our prediction about increasing temperature favoring the slow grower holds in the presence of two resources, however, is a relevant question in view of our experiments, in which we used a growth medium with two carbon sources (see Methods).

The LRC model with two resources can be solved analytically with a graphical method that we will not describe in detail here. It is sufficient to say that there is a region in the space of the two resources where both species can stably coexist, if the resource supply point lies in this region. In this case, an expression for the stable relative fraction of the slow grower can be obtained. We investigated whether this fraction increases with increasing temperature, and found that it does not always increase. The LRC model with two resources therefore makes no prediction about the effect of an increasing temperature on the slow grower.

The Monod model with two resources can be visualized with the same graphical method, but cannot be solved analytically. We relied on simulations to determine whether the relative fraction of the slow grower increases with increasing temperature, and found that this is not always the case, meaning that neither two-resource model makes a simple prediction about the effect of an increasing temperature. Our findings are summarized in Supplementary Table 1.

| Model | Prediction |
|---|---|
| Lotka-Volterra | * |
| Linear Resource Concentration, one resource | * |
| Monod, one resource | ** |
| Linear Resource Concentration, two resources | -- |
| Monod, two resources | -- |

**Supplementary Table 1**: Predictions about the effect of increasing temperature are summarized. One asterisk means that the model always predicts that the slow grower is favored by increasing temperature, assuming that it does not become relatively faster at high temperature. Two asterisks mean that the same prediction is true, with the additional assumption that the slow grower does not become relatively faster at low resource concentration. A double dash means that the model makes no simple prediction about the effect of an increasing temperature.
